# Distinct mitochondrial remodeling during early cardiomyocyte development in a human-based stem cell model

**DOI:** 10.1101/2021.07.07.451436

**Authors:** Sepideh Mostafavi, Novin Balafkan, Ina Katrine Nitschke Pettersen, Gonzalo S. Nido, Richard Siller, Charalampos Tzoulis, Gareth Sullivan, Laurence A. Bindoff

**Affiliations:** Department of Clinical Medicine, University of Bergen, Norway; Department of Mental Health Research, Haukeland University Hospital, Norway; Norwegian Centre for Mental Disorders Research (NORMENT) – Centre of Excellence, University of Oslo, Norway; Institute for Biomedicine, University of Bergen, Norway; Stem Cell Epigenetics Laboratory, Institute of Basic Medical Sciences, University of Oslo, Norway; Neuro-SysMed, Center of Excellence for Clinical Research in Neurological Diseases, Department of Neurology, Haukeland University Hospital, Norway; Department of Molecular Medicine, Institute of Basic Medical Sciences, University of Oslo, Norway; Norwegian Center for Stem Cell Research, Oslo University Hospital and the University of Oslo, Norway; Institute of Immunology, Oslo University Hospital, Norway; Hybrid Technology Hub – Centre of Excellence, Institute of Basic Medical Sciences, University of Oslo, Norway; Department of Pediatric Research, Oslo University Hospital, Norway

**Keywords:** Mitochondria, mtDNA, hPSC, hiPSC, OXPHOS, cardiomyocyte differentiation, development

## Abstract

Given the considerable interest in using stem cells for modelling and treating disease, it is essential to understand what regulates self-renewal and differentiation. Remodeling of mitochondria and metabolism, with the shift from glycolysis to oxidative phosphorylation (OXPHOS), play a fundamental role in maintaining pluripotency and stem cell fate. It has been suggested that metabolic ‘switch’ from glycolysis to OXPHOS is germ-layer specific as during early ectoderm commitment, glycolysis remains active while during the transition to mesoderm and endoderm lineages, it is downregulated. How mitochondria adapt during these metabolic changes and whether mitochondria remodeling is tissue specific remains unclear. Here we address the question of mitochondrial adaption by examining the differentiation of human pluripotent stem cells to cardiac progenitors and further to functional cardiomyocytes. Contrary to recent findings in neuronal differentiation, we found that mitochondrial content decreases continuously during mesoderm differentiation, despite clear mitochondrial remodeling giving increased mitochondrial activity and higher levels of ATP-linked respiration. Thus, our work both highlights similarities in mitochondrial remodeling during the transition from pluripotent to multipotent state in ectodermal and mesodermal lineages, while at the same time demonstrating cell-lineage-specific adaptions upon further differentiation. Our results improve understanding of how mitochondria remodeling and the metabolism interact during differentiation and show that it is erroneous to assume that increased OXPHOS activity during differentiation requires a simultaneous expansion of mitochondrial content.

**Summary statement:** We found that mitochondrial content decreases continuously during mesoderm differentiation, despite clear mitochondrial remodeling giving increased mitochondrial activity and higher levels of ATP-linked respiration during mesoderm differentiation.

## Introduction

Crosstalk between mitochondria, metabolism and processes controlling stem cell fate is essential (reviewed in(Wanet et al., 2015)). As human pluripotent stem cells (hPSCs) exit pluripotency they switch from a primarily glycolytic based metabolism that provides the energy and substrates necessary for proliferation in a hypoxic niche, to one more dependent OXPHOS, which is better suited for post-mitotic tissues with high energy demand(Locasale and Cantley, 2011; Varum et al., 2011; Zhang et al., 2012a). Metabolic switching during differentiation is thought to be germ layer specific, as inhibition of glycolysis inhibits neuronal differentiation, but has no effect on mesodermal and endodermal differentiation(Cliff et al., 2017). Also, the MYC transcription factor family that drives glycolysis remains activated after exit from pluripotency in nascent ectoderm, while it is silenced during mesoderm and endoderm differentiation(Cliff et al., 2017). How mitochondrial properties are adapted through metabolic switching during early stages of differentiation across different germ-layers remains unclear.

It has been proposed that the switch from glycolysis to OXPHOS is the result of remodeling mitochondria from a fragmented state with lower mitochondrial DNA (mtDNA) and mass in hPSCs, to elongated mitochondria with high levels of mtDNA and mitochondrial mass in differentiated cells(Sercel et al., 2021). Experimental data for this hypothesis is, however, limited either to what happens in neuronal differentiation(Zheng et al., 2016; Lees et al., 2018) or to earlier studies describing mitochondrial changes during mesoderm and endoderm differentiation from heterogeneous cultures derived from embryoid bodies(John et al., 2005; Cho et al., 2006; Belmonte and Morad, 2008; Lukyanenko et al., 2009). Noteworthy, ectoderm differentiation showed a biphasic change in mitochondrial content: in early differentiation, mitochondrial mass, mtDNA and superoxide production decreased while in the later stages, there was an expansion of mitochondrial mass and mtDNA along with a higher OXPHOS activity(John et al., 2005; Cho et al., 2006; Zheng et al., 2016; Lees et al., 2018).

In order to gain greater insight into the metabolic switch during mesoderm differentiation, we assessed mitochondrial properties including mitochondrial abundance, ultrastructure, membrane potential, and respiratory complex activity during differentiation and maturation of cardiomyocytes derived from both human embryonic stem cells (hESC) and human induced pluripotent stem cells (hiPSC). Cardiomyocytes derived from hPSC share the characteristics and functional properties of primary human heart tissue and are able to recapitulate the in vivo developmental process(Zwi et al., 2009; Zhu et al., 2017; Friedman et al., 2018).

In contrast to previous reports, we detected a significant reduction in mitochondrial biomass and mtDNA levels during differentiation of hPSC towards cardiac progenitors and functional cardiomyocytes. Despite this marked mitochondrial reduction, however, differentiated cells showed a higher mitochondrial coupling efficiency and appeared more dependent on OXPHOS with a higher mitochondrial membrane potential per unit mitochondria than undifferentiated hPSCs. Overall, our findings suggest a unique mitochondrial remodeling process for cardiomyocyte differentiation whereby mitochondrial biogenesis decreases during the transition from hPSC into differentiated cardiomyocytes, while the efficiency of ATP generation through OXPHOS increases in keeping with mitochondrial maturation.

## RESULTS AND DISCUSSION

### Cardiomyocyte differentiation and characterization

Two hESC lines (429 and 360) and three hiPSC lines (established from two independent Detroit 551 clones (clones 7 and 10), one CRL 2097 iPSC clone, CRL-8) were selected for cardiomyocyte differentiation in a 96-well plate format(Balafkan et al., 2020). For comparative purposes, we divided the differentiation process into phases based on the expression of cell-type-specific markers (Fig. 1A): pluripotent state (S1, day 0), mesendoderm cells (S2, day 1-2), cardiac mesoderm (S3, day 3), cardiac progenitor cells (S4 - day 5-7), functional cardiomyocyte (S5, day 12-15). The specific markers for the mesoderm cell lineage were assigned based on previous studies(Sturzu and Wu, 2011; Vliet et al., 2012). We have previously shown that cardiomyocytes differentiated using this protocol in a 96-well plate format are physiologically functional as determined by assessing different ion channels and beat interval (see Movie 1-2)(Balafkan et al., 2020). Initial transcriptomic profiling of cardiac progenitors derived from different hPSCs was performed with samples collected from four hiPSCs and two of the hESC lines (429 and 360) at pluripotent state (S1) and cardiac progenitor (S4) (Fig S1). In addition, to validate our gene expression analysis and further profile the transcriptomes of the critical stages of mesoderm differentiation including S5, an independent RNA-seq experiment was performed using a hESC line (H1), which solely used for transcriptomic analysis and not included in other set of experiments. RNA samples from three independent differentiations of H1 were collected at the following stages: pluripotent state (S1), mesendoderm state (S2), cardiac progenitor (S4) and functional cardiomyocyte (S5). Both data sets revealed downregulation of pluripotent stem cell markers, while up-regulated genes were enriched in pathways associated with cardiomyocyte differentiation including regulation of cardiac muscle, ventricular cardiac muscle tissue morphogenesis, sarcomere, and regulation of heart contraction (Fig. 1B and S1). The correct cardiomyocyte differentiation route was confirmed using qPCR (Fig. S1) and immunocytochemistry (Fig. 1C and 1D). This data indicates that our model is a reliable model to assess cardiomyocyte differentiation.

**Figure 1).**
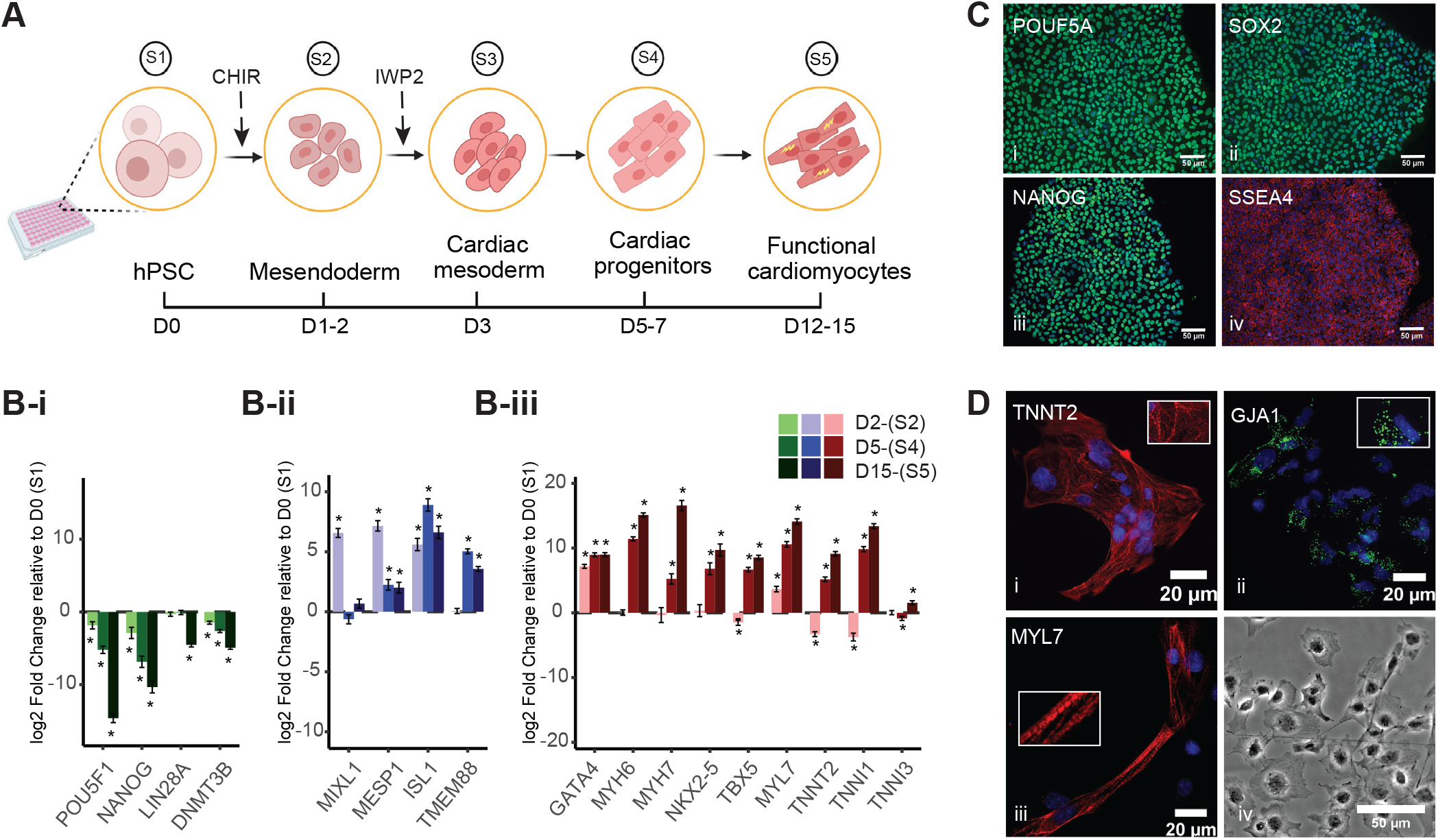
Cardiomyocyte differentiation and characterization. A) Schematic diagram of the cardiomyocyte differentiation protocol. Cells were treated with 6 *μ*M (hiPSCs) or 8 *μ*M (hESCs) CHIR99021 (CHIR) on day 1 for 24 h followed by treatment with IWP2, a Wnt signaling inhibitor, for 48 h on day 3. We divided cardiomyocyte differentiation into five different stages (S1-S5) for comparative purposes. B) Relative expression of known markers for stem cells (B-i), cardiac progenitor (B-ii) and cardiomyocytes (B-iii) haven been assessed using RNAseq data from H1 (n=3). C and D) Both hPSCs and hPSC-derived cardiomyocytes examined for expression of specific markers using a battery of antibodies. Pictures show co-immunostaining of the cell specific markers (labelled on the image) and nucleus (blue), E-iv represents a bright field image of cardiac cells at S5 (transferred from a 96-well plate to cover glass at S4). Scale bars demonstrates 50*μ*m, otherwise are mentioned on the picture.

Previously, we have shown that differentiating cells in a 96-well plate format results in a population of both cardiomyocytes and non-cardiomyocytes, similar to what is expected in the composition of the mammalian heart(Sturzu and Wu, 2011). The TNNT2+ cells exhibit relevant electrophysiological properties and express specific cardiomyocyte markers, while the non-cardiac population consists of different mesodermal lineages such as smooth muscle and endothelial cells (Fig. S2)(Skelton et al., 2017; Balafkan et al., 2020). Thus, cells in S1 represent a pure population of PSCs (SSEA4+), S4 a relatively pure population of cardiac progenitors (ISL1+), while S5 comprises two different cell populations, TNNT2+ and TNNT2-cells, i.e. cardiomyocytes and non-cardiomyocyte cell populations (Fig. S2). The presence of both cardiomyocytes and non-cardiomyocytes is a particularly important aspect of our study since we have, for the first time, investigated the differences between two major cell subpopulations resulting from mesoderm differentiation.

Of note, since there were no significant differences in mitochondrial properties between hESCs and hiPSCs during cardiac differentiation, we pooled the results from the hiPSC and hESC lines for each stage. We demonstrated inter- and intra-individual variation for each set of experiments in supplementary data.

### Mitochondrial content falls progressively during mesoderm differentiation

Thirteen polypeptides that are essential respiratory chain components are encoded by mtDNA. Unlike nuclear DNA, mtDNA is present in multiple copies and its copy number can impact the levels of mitochondrial RNA transcripts available for generating respiratory chain subunits(Anderson et al., 1981). We assessed the mtDNA copy number at different stages of cardiomyocyte differentiation using real-time PCR quantification relative to the nuclear gene APP(Tzoulis et al., 2013). This revealed a clear and progressive reduction of mtDNA copy number (of up to 85%) during differentiation of both hiPSC and hESC to cardiomyocyte lineage (Fig. 2A). Others have found that mtDNA levels drop during ectoderm differentiation and reach their lowest levels in NPCs(Lees et al., 2018). Our findings showing low mtDNA level in cardiac progenitors (S4) is, therefore, similar to what has been reported in ectoderm differentiation(Lees et al., 2018), however, fact that the mtDNA levels did not rise in S5 relative to PSC was unexpected. More recent findings showed a higher level of mtDNA and mitochondria content in fully differentiated cells relative to undifferentiated cells derived from hPSC, which is aligned with the transition from glycolytic state to a state with higher OXPHOS activity(Malinska et al., 2012; Wanet et al., 2014; Zheng et al., 2016). For comparative purposes, we quantified mtDNA in post-mortem human heart using the same method and confirmed that the level of mtDNA in the mature tissue is at least 11-44 times higher than hPSCs and differentiated cells at S5 (Fig. 2B).

**Figure 2).**
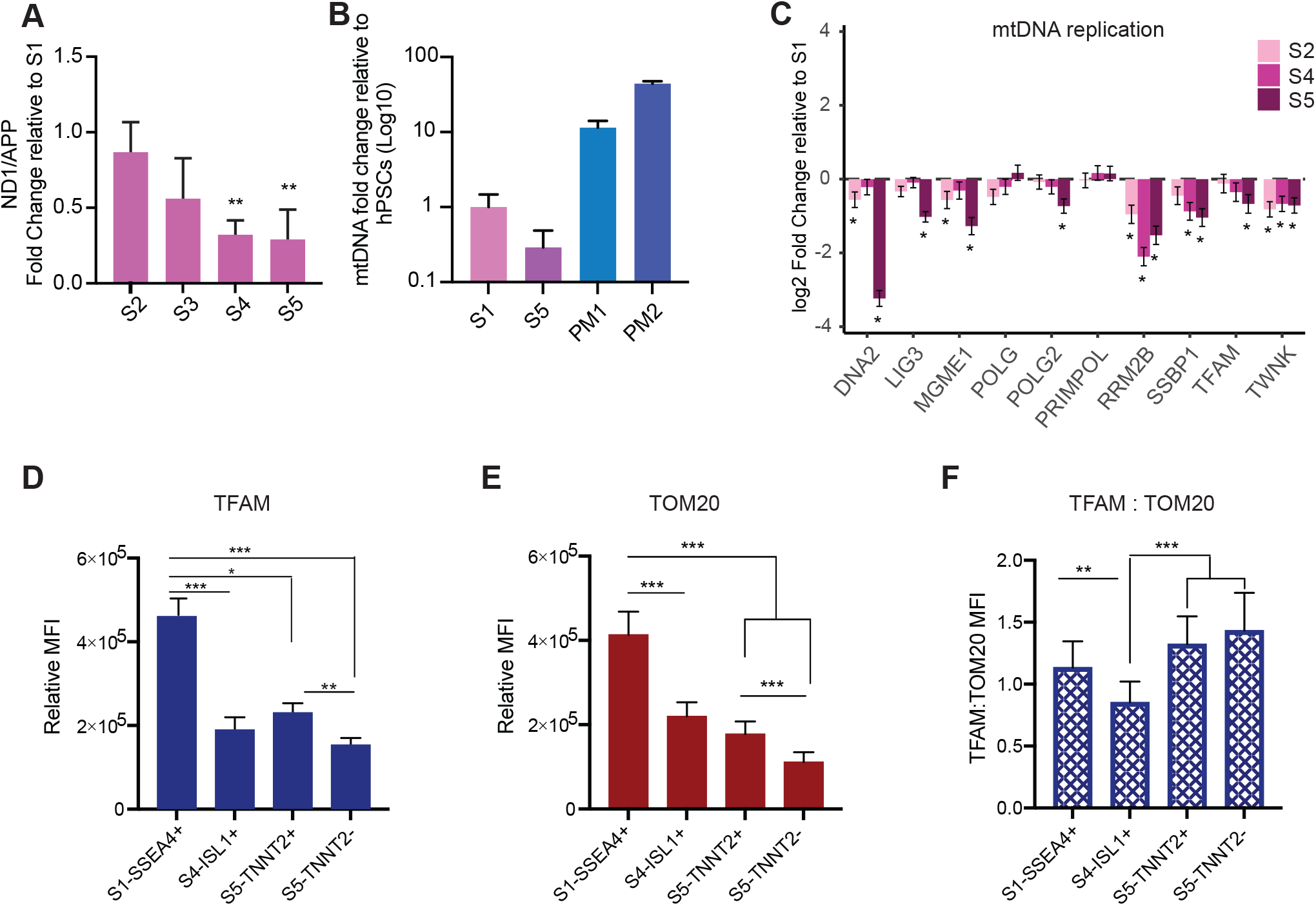
Changes in mitochondrial mass and mtDNA during cardiomyocyte differentiation. **A)** MtDNA copy number fell progressively during cardiomyocytes differentiation. Each bar represents the mean of the mtDNA fold change, pooled data from both hESCs and hiPSCs, at each stage relative to undifferentiated cells at S1 and error bars represent mean values ± 95% c.i., n=11. Statistical significance was assessed using one-way ANOVA followed by Dunnett’s post-hoc multiple comparisons. **B)** mtDNA levels were significantly lower in both S1 and S5 cells compared to human heart tissue. The bar chart demonstrates the average mtDNA level in collected cells from all 5 hiPSC lines at S1 (n=9) and S5 (n=8) in comparison with mtDNA level of cells isolated from post-mortem heart tissues (PM1: 59y, female, died of lung cancer, and PM2: 65y, male, died of colon cancer). **C)** Changes in transcriptional level of a battery of genes involved in mtDNA replication and homeostasis at different stages of differentiation relative to undifferentiated stage (S1) in H1 (n=3). **D)** We measured TFAM levels using flow cytometry and detected a significant loss of TFAM at stage S4 and S5 compared with the level at S1. Also, there was a significant difference in TFAM level between TNNT2+ and TNNT2-cells. **E)** We measured TOM20 using flow cytometry and a similar significant decrease at S4 and further reduction was recorded at S5. TNNT2+ demonstrated significantly higher level of TOM20 relative to TNNT2-cells. **F)** The ratio of TFAM:TOM20 increased significantly from stage S4 to stage S5 of cardiac differentiation. Since TFAM binds mtDNA in stoichiometric amounts, this suggests that mtDNA levels have increased per unit mitochondrial volume. In bar charts D-E; each bar represents the mean of the relative median fluorescent intensity (MFI)±s.d., n=18 (SSEA4+ and ISL1+), 20 (TNNT2+/-), and * P < 0.05, ** P < 0.01 and *** P < 0.001.

Assessed by bulk RNA-seq, we found that the majority of genes involved in mtDNA homeostasis decreased during mesoderm differentiation, including mtDNA maintenance exonuclease 1 (MGME1), single-stranded DNA binding protein (SSBP), mitochondrial transcription factor A (TFAM), and mtDNA helicase (TWNK) (Fig. 2C and Fig. S3). We validated RNA-seq findings by using real-time PCR to quantify the expression of SSBP and *POLG* that both have critical roles in mtDNA replication and were able to show a progressive decline of SSBP with no significant change in POLG expression level (Fig. S3). No evidence of qualitative damage such as mtDNA deletion was found at any stage of differentiation (Fig. S3F and G).

The link between the TFAM and mtDNA has been investigated in depth(Larsson et al., 1994, 1998) and studies show that it binds mtDNA in molar quantities (Ekstrand et al., 2004; Kaufman et al., 2007; Kukat and Larsson, 2013). Using flow cytometry, we assessed TFAM both as a direct measure of a mitochondrial matrix protein and as an indirect measure of mtDNA level within each cell population co-stained with antibodies against stage-specific markers to validate our findings from real-time PCR. We found a 58% reduction of TFAM in cardiac progenitors (S4) relative to the pluripotent stage (S1) (Fig. 2D). Next, we examined changes in mitochondrial mass during cardiomyocyte differentiation at the single-cell level. We used flow cytometry and co-staining with antibodies against stage-specific markers and Translocase of Outer Mitochondrial Membrane 20 (TOMM20), a well-established marker of mitochondrial mass. The TOMM20 level showed a significant fall (43%) from S1 to S4 and reached its lowest level at S5 (60%) (Fig. 2E). As expected, the level of TOM20 was lower (∼40%) in non-cardiomyocytes than cardiomyocytes (TNNT2+) indicating a lower level of mitochondria in non-cardiac cells. The results from each line for each stage of differentiation are shown in figure S4. The fall in mitochondrial mass starting from S3 was confirmed by a significant reduction of VDAC gene expression; VDAC encodes a vital outer membrane protein that is also routinely used as a mitochondrial mass marker (Fig. S3E). Given the link between the cell size and mitochondrial mass(Zheng et al., 2016), the finding of a lower mitochondrial mass was surprising; differentiated cardiomyocytes are much larger cells than PSC (Fig. 1D-iv). A reduction of both mtDNA and mitochondria mass was previously found in the very early stage of ectoderm differentiation, but both started to increase from NPC onward(Birket et al., 2011; Zheng et al., 2016; Lees et al., 2018). A similar decline in mtDNA content has also been reported during hematopoietic differentiation as well as during differentiation of hESC toward primordial germ cells (Almeida et al., 2017; Floros et al., 2018). Thus, our findings in cardiac progenitors suggest that the reduction of mtDNA levels and mitochondria content from pluripotent to multipotent stem cells may be a more general phenomenon than previously thought, and not limited to one specific cell lineage.

Earlier studies have shown that the undifferentiated cells at the embryoblast stage (i.e., the inner cell mass) have the lowest level of mtDNA level and mitochondria content(Taylor and Pikó, 1995; Floros et al., 2018), and spontaneous differentiation through formation of embryoid bodies resulted in an increase in mtDNA proliferation and mitochondria mass(John et al., 2005; Cho et al., 2006; Belmonte and Morad, 2008; Lukyanenko et al., 2009). The reduction of mtDNA and mitochondrial mass in early ectoderm differentiation partially can be explained by the fact that ectodermal progenitors are less relay on OXPHOS, while cells at early stages of mesoderm differentiation are heavily relay on OXPHOS(Cliff et al., 2017). Another possible explanation is that this reduction in mitochondrial content seems to set the scene for a mitochondrial bottleneck to select mtDNA with specific variants and mitochondria specialized with a certain mode of energy production required for development of specific germ-layers(Gong et al., 2015; Floros et al., 2018; Rossmann et al., 2021).

Interestingly, when we measured changes in mtDNA level relative to mitochondrial content, by plotting the level of TFAM (an indirect marker of mtDNA) against TOM20 (a direct marker for mitochondrial content), we found that this ratio varied during the course of cardiomyocyte differentiation. The ratio of TFAM to TOM20 was significantly higher in differentiated cells in S5 relative to cardiac progenitors (S4) (Fig. 2F) suggesting that the lowest level of mtDNA per unit mitochondria mass occurred in cardiac progenitors and then rose in the more differentiated stage. These findings also highlight that mtDNA levels may not act as a proxy for mitochondrial level, particularly during differentiation, and it is important to use a complementary method to assess mitochondrial levels directly.

### Despite lower mitochondrial content, differentiated cells generate more energy through TCA than glycolysis

Despite analyzing a heterogeneous population (TNNT2+ and TNNT2-cells), we were able to verify a significant increased expression of mtDNA genes in the S5 stage relative to other stages of differentiation (Fig. 3A) via bulk RNA-seq. These observations suggest that differentiated cells are relying more on mitochondrial respiration, regardless of the cell type they are committed to, and despite having much lower mitochondria mass.

**Figure 3).**
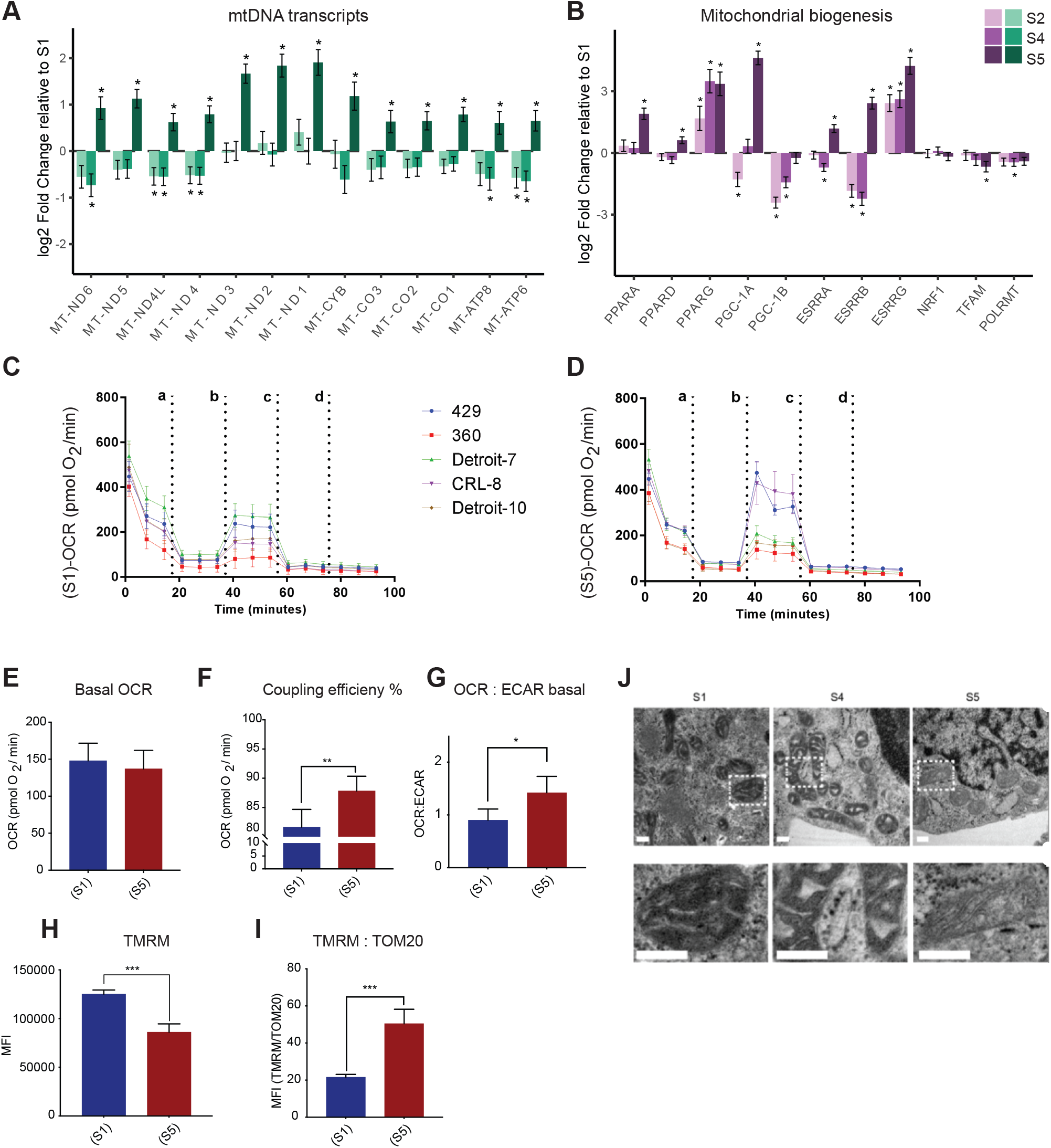
Mitochondria respiration in hPSCs and differentiated mesodermal derivatives. **A)** The mtDNA transcripts collected via RNA-seq at different stages of cardiomyocyte differentiation suggest a significant increase in the number of mitochondrial transcripts in S5 relative to S1, S2 and S4. **B)** A battery of genes associated with metabolic remodeling selected based on previous studies (Scarpulla et al., 2012; Zheng et al., 2016) to evaluate their gene expression changes at different stages of cardiomyocyte differentiation relative to undifferentiated stage S1. Barplots depict the estimated log 2 fold change±s.e.m, n=3. **C and D)** Graphs present the result of oxygen consumption rate (OCR) in hPSCs and differentiated cells at S5. Cells were exposed sequentially to oligomycin (a), CCCP (b), rotenone (c) and antimycin A (d). **E)** To better visualize the result of Seahorse we presented data as bar charts. Basal OCR level didn’t change in differentiated cells relative to undifferentiated hPSCs, which suggest higher level of oxygen consumption by differentiated cells since we have shown these cells have almost 50% less mitochondria compare to hPSCs. **F)** Coupling efficiency significantly increased in differentiated cells, which shows the higher potential for ATP generation in differentiated cells relative to undifferentiated hPSCs. **G)** The basal OCR/ECAR showed an increase in differentiated cells, which shows these cells are more relay on OXPHOS compare to undifferentiated hPSCs. **H)** We found a high concentration of TMRM in hPSCs relative to S5 cells, however after correction for mitochondrial content in differentiated cells we found (I) a significant increase of TMRM-MFI level in S5 cells relative to undifferentiated hPSCs. **I)** The ratio of TMRM and TOM20 suggested higher level of MMP per unit of mitochondrial mass of differentiated cells compared with undifferentiated hPSCs. Each bar shows mean±s.d., n=3 or 4 independent experiments, and t-test was chosen to assess the statistical significance of the difference between undifferentiated hPSCs and S5. In all bar charts; * P < 0.05, ** P < 0.01 and *** P < 0.001. **J)** Representative TEM pictures of mitochondria during cardiomyocyte differentiation of a hiPSC line (Detroit) at S1, S4 and S5. Mitochondria appear smaller with wider more immature cristae at stages S1 and S4, while at S5, more mitochondria show an expanded matrix with typical, narrow mature cristae. The bottom panel is the magnification of the indicated areas in the upper panel. Scale bar: 200 nm.

We then assessed mitochondrial oxidative activity by measuring oxygen consumption rate (OCR) in differentiated cells at S5, which contain beating cardiomyocytes (see Movie 3-4), and in undifferentiated stem cells (S1) using Seahorse XF-96 extracellular flux analyzer (Fig 3C, 3D and Fig. S5). Interestingly, despite a major drop in mitochondrial mass, no major change in basal respiration, the amount of oxygen consumed by mitochondria under basal conditions, was detected: differentiated cells at S5 137 pmol/min, undifferentiated hPSCs at S1 148 pmol/min (Fig. 3E). Maximal OCR and spare capacity (also known as reserve capacity) showed a slight increase in S5 compared to S1 cells (Fig. S5B and S5C). Coupling efficiency, the proportion of mitochondrial respiratory chain activity used to drive ATP synthesis was significantly higher (Fig 3F)(Divakaruni and Brand, 2011). The ratio of OCR to extracellular acidification rate (ECAR), an indicator of how much lactate is produced through glycolysis also rose in S5 cells relative to S1 suggesting a shift to mitochondrial respiration (Fig 3G). These findings corroborate previous studies showing the change in energy profile between PSC and terminally differentiated cells(Folmes et al., 2012). The finding of a negative value for spare capacity in hPSC lines (360 and CRL-8), raised the possibility that uncoupling by CCCP had collapsed the membrane potential in these lines (see blue bars in Fig. S5A-iv), and that, with a respiratory chain already working maximally, there was no further reserve capacity to be used with the extra stress(Varum et al., 2011; Zhang et al., 2012b).

We also investigated mitochondrial membrane potential (MMP) as a marker for mitochondrial activity. Membrane potential is generated by proton movement driven by the ETC and although MMP is an excellent marker for assessing mitochondrial function, it also reflects mitochondrial volume(Perry et al., 2011). Therefore, we employed a lipophilic cationic fluorescent probe tetramethylrhodamine methyl ester (TMRM) to evaluate MMP at the single-cell level by flow cytometry and normalized our data to mitochondria mass measured via TOM20 in the previous step. The absolute median fluorescent intensity (MFI) of TMRM was significantly lower at S5 relative to S1 (Fig. 3H and Fig. S5E), however, when we adjusted the MFI of TMRM for the mitochondrial mass, using the ratio of MFI-TMRM to MFI-TOM20, we found a more than two-fold increase in MMP at S5 compared with undifferentiated hPSCs at S1 (Fig. 3I and Fig. S5F). The significantly higher level of MMP per unit of mitochondrial mass in differentiated cells at S5 relative to undifferentiated hPSCs was further proof that the mitochondria in differentiated cells are more efficient in generating ATP.

### Cristae remodeling during cardiomyocytes differentiation

We identified a significant increase in the expression of genes associated with mitochondrial biogenesis and respiration activities (e.g., PPARA, PPARG, PGC-1A and ESSRA, see figure 3B): since no increase in mitochondria mass was demonstrated (Fig. 2E), this suggested remodeling of mitochondrial membrane and cristae. We examined cells collected on S1, S4 and S5 by transmission electron microscopy (TEM) in order to assess the mitochondrial morphology and cristae structure. We observed that mitochondria went from having wide cristae with a dense matrix to having more compact cristae and clearer matrix (Fig. 3J). Previous findings have revealed that mitochondrial inner membrane morphology modulates OXPHOS function by modifying the kinetics of chemical reactions and regulating the arrangement of protein complexes(Cogliati et al., 2016). It is also known that modulating cristae structure affects the mitochondrial respiratory efficiency independently of changes to mitochondrial protein synthesis(Cogliati et al., 2013). Our findings align with previous studies that reported the formation of cristae and mitochondrial permeability transition pore (mPTP) closure in differentiated cardiomyocytes, which are known characteristics of the developmental maturation of mitochondria(Hom et al., 2011; Teixeira et al., 2015; Dai et al., 2017).

Our work highlights similarities in mitochondrial remodeling in the transition from pluripotent to multipotent state in ectodermal and mesodermal lineages, while at the same time suggesting that there are cell-lineage-specific metabolic paths following commitment to a specific cell type. This is in contrast to earlier studies that suggested a simultaneous expansion of mitochondrial content and increase in OXPHOS activity during all lineage differentiation. Interestingly, our findings also offer experimental validation for the previously proposed mathematical model that suggested cells modulate their mitochondrial membrane potential, rather than their mtDNA level, to quickly meet the changes in energy demands in mammalian cells(Miettinen and Björklund, 2016; Aryaman et al., 2017). More importantly, our findings emphasise the importance of differentiating between active loss of mtDNA and failure to expand mtDNA copy number when studying tissues derived from mesodermal lineages in infants with mitochondrial disease.

Whether our findings are physiologically relevant or reflect changes in signaling pathways during the early stages of the differentiation protocol under the normoxic conditions, remains an unanswered question. Nevertheless, our findings do emphasise the importance of understanding how mitochondrial properties change across different cell lineages and highlight the narrow time window that is crucial in the relationship between mitochondria remodeling and cell fate that can be a target for future investigations.

## ACKNOWLEDGMENTS

The authors wish to acknowledge the help and resources provided by the Flow Cytometry Core Facility, University of Bergen and the Molecular Imaging Center (MIC), Department of Biomedicine, University of Bergen. We would like to thank Tilo W. Eichler and Christian Dölle for helpful discussions. This work was supported by grants from The Research Council of Norway (NFR, project no. 229652), Bergen Stem Cell Consortium (BSCC), Haukeland University Hospital, University of Bergen and the Rakel og Otto-Kristian Bruun’s Legat.

## AUTHOR CONTRIBUTIONS

**Conceptualization**: L.A.B, N.B.; **Methodology**: N.B., S.M., G.S.; **Software**: G.S.N.; **Validation and formal analysis**: N.B., S.M., G.S.N.; **Investigation**: S.M., N.B., I.K.N.P, R.S.; **Resources**: L.A.B, G.S., C.T.; **Data Curation**: G.S.N., N.B.; **Writing – original draft preparation:** N.B.; **Writing – review and editing**: L.A.B, N.B., S.M.; **Visualization**: S.M., N.B., G.S.N.; **Supervision**: L.A.B, N.B.; **Project administration and funding acquisition**: L.A.B.

## MATERIALS AND METHODS

### Cell lines

Three hESC lines (H1, 429 and 360) and four hiPSC lines (established from two independent Detroit 551 clones (clones 7 and 10), one CRL 2097 iPSC clone, CRL-8) were employed for this study. The details of reprogramming and characterization of the lines hiPSC published elsewhere(Balafkan et al., 2020; Liang et al., 2020). Passage number of the lines were kept between 23-55 for all the experiments.

### Cardiomyocyte differentiation

Cardiomyocyte differentiation was performed in 96-well microplate as previously described(Balafkan et al., 2020). Briefly, hiPSCs were seeded at 2.4 × 10^4^ cells/cm and propagated on Geltrex (#A1413302, Thermo Fisher Scientific) diluted (1:100) in Advanced DMEM/F-12 (#12634010, Thermo Fisher Scientific), under feeder free conditions in Essential 8 Medium (E8) (#A1517001, Thermo Fisher Scientific). Within 3 days, when cells reached the optimum confluency, 60 to 70%, cardiomyocyte differentiation was started by applying the GSK3 inhibitor CHIR99021 (#4423, Tocris Bioscience) in RPMI 1640 (#61870, Thermo Fisher Scientific) medium supplemented with B27-without insulin (RPMI-B27) (#A1895601, Thermo Fisher Scientific), in concentration-cell-dependent manner. After 24h, medium was changed to RPMI-B27 without CHIR99021. Differentiation process was continued by adding 5 *μ*M inhibitor of WNT production-2, IWP2, (#3533, Tocris Bioscience) diluted in RPMI-B27, 72 hrs post differentiation induction for 48 hrs. Fresh RPMI-B27 medium was provided on day 5, and from day 7 cells were fed with fresh RPMI medium supplemented with B27 with insulin (#17504044, Thermo Fisher Scientific) without extra supplement) every two other days.

### Gene expression analysis

MagMAX™-96 Total RNA Isolation Kit (#AM1830, Thermo Fisher Scientific) was used for RNA isolation from cultured cells. Cells were rinsed with DPBS (#14190250, ThermoFisher) and lysis buffer provided in the kit was added directly to the cells and lysates were either immediately used for RNA isolation using automated MagMAX express 96 or stored at −80°C. EXPRESS One-Step Superscript qRT-PCR Kit (#11781-01K, Thermo Fisher Scientific) was used for cDNA synthesis and real-time PCR using TaqMan probes (Table S1) on an Applied Biosystems 7500-Fast real-time PCR System (Thermo Fisher Scientific). All real-time PCR reactions were performed in triplicate and the average of Ct values normalized to the geometric mean of ACTB and GAPDH as endogenous control genes, and the result (dCt) used for further analysis. Data was represented as the mean of 1/dCt with 95% CI of the mean and significance level of difference in expression of each gene at different time points was calculated using one-way ANOVA, whereas the difference between hiPSC and hESC lines at each time point was calculated using multiple t-test with Holm-Sidak correction for multiple comparison.

### RNA sequencing

RNA sequencing was carried out on two different data sets, which analyzed independently and separately. Data set A contains RNA sequences from both hESCs and hiPSC lines collected from S1(Undifferentiated cells-D0) and S4 (Cardiac-progenitors-D7) of cardiomyocyte differentiation. It composed of samples collected from three independent differentiation of two hESC lines (429 and 360), two independent differentiation of Detroit-7 and CRL-8, and one differentiation of Detroit-10. Data set B constitutes from RNA sequences collected from four stages S1-S5 (D0, D2, D5, and D15) of three independent differentiation runs of the H1 line (hESC). The total RNA samples were isolated using MagMAX™-96 Total RNA Isolation Kit.

RNA sequencing for the dataset (A) was carried out at the Finnish Microarray and Sequencing Centre’s analysis service and Biocenter Finland. The quality of the total RNA samples and libraries was ensured with Advanced Analytical Fragment Analyzer. Sample concentration was measured with Qubit® Fluorometric Quantitation, Life Technologies. RNA quality as measured by RNA integrity number (RIN) was well above 7 for all samples (median RIN = 9,25) with the exception of a single sample, with RIN = 5.5. The samples were sequenced with Illumina HiSeq 3000 instrument. Single-read sequencing with 1 × 50 bp read length was used, followed by dual index 8 bp run. The base calling was performed using Illumina’s standard bcl2fastq2 software, automatic adapter trimming was used. RNA sequencing for the second dataset (B) was carried out at the HudsonAlpha Genome Sequencing Center, USA. RNA quality as measured by RNA integrity number (RIN) was well above 7 for all samples (median RIN = 8,8).

### RNA expression quantification and filtering

RNA sequencing analyses were performed independently for each of the datasets. Raw FASTQ files were assessed using fastQC version 0.11.8 (Andrews, 2010), and reads were mapped using Salmon version 1.3.0 (Patro et al., 2017) with the fragment-level GC bias correction option (-gcBias) against the GENCODE Release 32 (GRCh38.p13) reference transcriptome and the GRCh38 reference genome, including the whole genome as decoy sequences. Transcript quantification was collapsed to the gene-level using the R package tximport version 1.14.2 (Soneson et al., 2015) with default parameters and the GENCODE Release 32 (GRCh38.p13) annotation. Low-expressed genes (i.e. genes with less than 10 reads in more than 75% of the samples of the corresponding dataset) were filtered out, resulting in 19,273 genes in dataset A, of which 80% annotated as protein-coding, and 22,480 in dataset B (73% annotated as protein-coding).

### Differential gene expression and functional enrichment analyses

Differential gene expression analyses were performed using the DESeq2 R package version 1.26 (Love et al., 2014)with default parameters. Samples from dataset A originated from different cell lines, and this was accounted for explicitly in the model by incorporating it as a covariate into the statistical model. Multiple hypothesis testing was performed with the default automatic filtering of DESeq2 followed by false discovery rate (FDR) calculation by the Benjamini-Hochberg procedure. Analyses were carried out independently for the two datasets. Genes were scored according to their significance by transforming the p-values to account for direction of change. In summary, for each gene, the up-regulated score (*S*_up_) was calculated as *S*_up_ = 1–p/2 if LFC < 0, and p/2 if LFC≥0. The down-regulated score (*S*_down_) as *S*_down_ = 1–*S*_up_, where LFC corresponds to the log fold change and p to the nominal p-value of the gene. Genes were then tested for enrichment using alternatively log(*S*_up_) and log(*S*_down_) scores employing the gene score resampling method implemented in the ermineR package version 1.0.1, an R wrapper package for ermineJ (Gillis et al., 2010) with the complete Gene Ontology (GO) database annotation(Ashburner et al., 2000), the KEGG database (Kanehisa and Goto, 2000) and a curated version of mitocarta (Calvo et al., 2016; Gaare et al., 2018) to obtain lists of up- and down-regulated pathways for each cohort. The source code for the RNA sequencing analyses is available in the GitLab repository (https://git.app.uib.no/neuromics/cardiomyocytes-rna-seq) under the GPL public license v3.0.

### Flow cytometry

Cells were washed in DPBS (#14190250, ThermoFisher) three times and dissociated into single cells suspension using TrypLE™ Express Enzyme (#12604013, Thermo Fisher Scientific) for 10-20 minutes at 37 °C and were collected in the micro centrifuge tube. After a quick wash with DPBS cells were stained with Zombie Red™ Fixable Viability Kit (#423110, BioLegend) according to manufacturer’s instructions. Single cells were fixed with 4% (vol/vol) paraformaldehyde for 10 min at room temperature (RT) and permeabilized for 10 min at −20°C with 90% ice-cold methanol diluted in PBS. Cells were blocked in blocking buffer consisting of 0.3 M glycine, 5% goat serum and 1% BSA in PBS for 20min at RT. The optimal concentration of conjugated antibodies for flow cytometry reported by the supplier for each batch was used. Antibodies are listed in Supplementary Table (Table S1). All the primary and secondary antibodies as well as conjugated antibodies diluted in blocking buffer and stained for 30 min at RT. Cells were washed after staining and collected in 300 *μ*l of 1% FBS in PBS. Samples were analysed quickly after staining. At least 30.000 events were collected for the target marker using a Sony cell sorter SH800 (Sony Biotechnology Inc.) and collected data analysed and presented by FlowJo V.10.5.0 (FlowJo LLC, OR, USA, www.FlowJo.com) 8-Peaks Rainbow Calibration Particles (Biolegend #422903) was used for routine alignment and performance verification of the flow cytometry. The flow cytometer was calibrated prior to quantitative fluorescence intensity measurements using Quantum™ Alexa Fluor® 488 MESF (molecules of equivalent soluble fluorophore) (Bangs Laboratories, Inc. #488). In order to test the integrity of the results over time, collected median fluorescence intensity (MFI) was normalized to MESF as an external control and the resulting values reported as relative MFI to MESF unit. All gates were adjusted according to fluorescence minus one control (FMO). To analyse the flow cytometry data the median values of MESF were used to compare the samples and 95% CI of the median used to show the confidence in median value.

### Immunocytochemistry and fluorescence microscopy

Cells were seeded either on Geltrex coated cover slips or in Millicell® EZ SLIDES (#PEZGS0816, Merck Millipore). Cells were fixed with 4% (vol/vol) paraformaldehyde for 10 min at RT and permeabilized with 0.3% Tween 20 (#822184, Merck Millipore). All the primary and secondary antibodies were diluted in blocking buffer consisting of 0.3 M glycine, 5% goat serum and 1% BSA in PBS. A final concentration of 10 *μ*g/ml was used for all primary antibodies and they were incubated overnight at 4°C. Alexa Fluor 488 or 594-conjugated (Thermo Fisher Scientific) secondary antibodies were diluted in blocking buffer 1:1000 and incubated for 30 min to 1h at room temperature. Nuclei were stained with Gold Antifade Reagent with DAPI (#P36935, Thermo Fisher Scientific). Confocal microscopy images were taken on eighter a Zeiss LSM 510 META or a Leica TCS SP5 at the Molecular Imaging Center (MIC), University of Bergen, and data analysis and image editing were done with Fiji (Schindelin et al., 2012). Antibodies are listed in Supplementary table (Table S1).

### Transmission Electron Microscopy

Cells were washed in DPBS and dissociated into single cells suspension using TrypLE™ Express Enzyme for 10-20 minutes at 37°C and were collected in the micro centrifuge tube and centrifuged at 300 g for 10 min at RT. Cells were fixed in 2,5% glutaraldehyde (diluted in a 0,1M sodium cacodylate buffer) for 24 hrs at 4°C and delivered to MIC facility at the University of Bergen. Post-fixation was performed for 1h (on ice) in 1% osmium tetroxide (EMS # 19134) diluted in 0,1 M sodium cacodylate buffer, followed by 2 washing steps. The samples were then dehydrated using a graded ethanol series (30%, 50%, 70%, 96% and 100%) before transferred to a 1:1 solution of 100% ethanol:propylene oxide (15 min). Samples were then transferred to 100% propylene oxide (15min) before gradually introducing agar 100 resin (AgarScientific R1031) drop by drop over the next hours. Samples were then transferred to a small drop of 100% resin and excess propylenoxid was allowed to evaporate (1h). Samples were then transferred to 100% resin and placed in molds and left in room temperature overnight. The molds were then placed at 60°C for 48h to polymerize. Ultrasections of approximately 60nm were placed on 100 mesh formvar coated(EMD # 15820) kopper grids (EMS #G100H-Cu) and stained with 2% uranyl acetate (EMS # 22400) and lead citrate (VWR #1.07398). Grids were imaged using a Jeol JEM-1230 transmission electron microscope at 80kV.

### MtDNA analysis

MtDNA quantification and deletion assessment was performed in DNA isolated from cultured cells using MagMAX™-96 DNA Multi-Sample Kit (#4413021, Thermo Fisher Scientific) and real-time PCR, as well as long range PCR, as previously described (Tzoulis et al., 2013). A commonly deleted region (MT-ND4) in the major arc of mitochondrial DNA and a rarely deleted region (MT-ND1) were utilized to quantify deletion and MT-ND1 was compared with amplification of a single-copy nuclear gene (APP) to assess the number of mtDNA copies. A triplex reaction of ND1, ND4 and APP was performed simultaneously within the same well using a 7500 fast sequence detection system (Thermo Fisher Scientific). Efficiency of the triplex reaction was measured prior sample analysis. Assessment of statistical significance was performed using one-way ANOVA with Dunnett’s post-hoc multiple comparisons.

### Measurement of OCR and ECAR using Seahorse XF-96 analyser

Respiration and acidification rates were measured on monolayer culture of undifferentiated hPSCs and cells at S5 using a Seahorse XFe96 extracellular flux analyser (Agilent, Santa Clara, CA, US). Geltrex coated XFe96 assay plates were utilized for seeding of 3×10^3^ undifferentiated hPSCs and 2×10^5^ hPSC-derived cardiac cells in each well. Cells were cultured in cell specific normal growth medium supplemented with 10 *μ*M of Y-27632 for 24 hours at 37 °C and 5% CO2). The next day, the supplemented growth medium was replaced with normal growth medium without supplement and kept till they reached almost 90% confluency prior to analysis. In order to find the optimal concentration of Carbonyl cyanide m-chlorophenyl hydrazone (CCCP) and oligomycin for each cell type, they were titrated prior to the cell analysis. The assay was performed in assay medium (pH 7.4, unbuffered) which was supplemented with 2 mM L-glutamine, 2 mM sodium pyruvate and 10 mM glucose. Cells were washed twice and pre-incubated in the assay medium and kept in a CO2-free incubator (XF Prep Station, Seahorse Biosciences) at 37°C for 1 hour before measurement, in order to remove CO2 from the medium. For mitochondrial respiration analysis, the final concentrations of 3 *μ*M oligomycin, 0.5-1 *μ*M CCCP, 1 *μ*M rotenone and 1 *μ*M antimycin A were diluted in assay medium. In order to correct the final results for differences in cell size and number between the undifferentiated hPSCs and differentiated cells at S5 we measured the total protein concentration for each well using absorbance at 280 nm. All the results were reported as pmol O2 per min after normalized to their total protein concentration. Each parameter was calculated accordingly; basal OCR= (OCR in non-treated cell – OCR after adding Antimycin A), Coupling Efficiency = (ATP Production Rate / Basal Respiration Rate × 100), spare respiration capacity = (maximal respiration / basal respiration). The XF reader software (Wave Desktop 2.4) was used to analyses the data.

### Measurement of mitochondrial membrane potential

We used a previously described protocol (Rowe and Boletta, 2013) to analyse the mitochondrial membrane potential (Ψm) independent of cell volume using TMRM. We quantified the fluorescent intensity of TMRM (FI) before and after applying Carbonyl cyanide 4-(trifluoromethoxy)phenylhydrazone (FCCP) as an uncoupler for OXPHOS which eliminates the mitochondrial membrane potential. The distribution of TMRM rely on the Nernstian equilibrium(Dykens and Stout, 2001). Accumulation of TMRM in the cytosol is very small compare with mitochondria due to the low negative charge of cytosol relative to mitochondrial matrix. Thus, fluorescence readout mirror the mitochondrial matrix:cell volume ratio. FCCP treatment of the cell releases the TMRM from mitochondrial matrix into cytosol and causes a re-equilibrium of TMRM in the cytosol. Therefore, fluorescence readout after FCCP treatment represents cell volume. The difference between geometric means for TMRM and TMRM-FCCP treated cells provided us the membrane potential level normalized to the cell size. We analysed mitochondrial membrane potential by flow cytometry starting with 1×10^6^ cells per sample. Each sample divided in two; one sample was treated with 25nM TMRM and another with a mixture of 25 nM TMRM and 100 *μ*M FCCP. Samples were incubated for 20 min at 37 °C and 5% CO2 washed three times with HBSS and analysed within 1 hour.

### Statistical tests

Data were expressed as mean or median with 95%CI of the mean or median depending on population distribution. To assess statistical significance, we used parametric tests; Student’s t test and ANOVA, and P values corrected with a multiple test correction by with Holm-Sidak and Dunnetts accordingly. The significance of median difference was assessed using non-parametric test; Kruskal-Wallis and Mann Whitney test followed by Dunn’s multiple correction. Data were analysed using GraphPad-Prism (Prism 7.0, GraphPad Software, La Jolla California USA).

## BIBLIOGRAPHY

Almeida, M. J. de, Luchsinger, L. L., Corrigan, D. J., Williams, L. J., and Snoeck, H.-W. (2017). Dye-Independent Methods Reveal Elevated Mitochondrial Mass in Hematopoietic Stem Cells. Cell Stem Cell 21, 725–729.e4. doi:10.1016/j.stem.2017.11.002.

Anderson, S., Bankier, A. T., Barrell, B. G., Bruijn, M.H.L., de Coulson, A. R., Drouin, J., et al. (1981). Sequence and organization of the human mitochondrial genome. Nature 290, 457–465. doi:10.1038/290457a0.

Andrews, S. (2010). FastQC: a quality control tool for high throughput sequence data. Available at: https://www.bioinformatics.babraham.ac.uk/projects/fastqc/RNA-Seq_fastqc.html.

Aryaman, J., Hoitzing, H., Burgstaller, J. P., Johnston, I. G., and Jones, N. S. (2017). Mitochondrial heterogeneity, metabolic scaling and cell death. Bioessays 39, 1700001. doi:10.1002/bies.201700001.

Ashburner, M., Ball, C. A., Blake, J. A., Botstein, D., Butler, H., Cherry, J. M., et al. (2000). Gene ontology: tool for the unification of biology. The Gene Ontology Consortium. Nat Genet 25, 25–9. doi:10.1038/75556.

Balafkan, N., Mostafavi, S., Schubert, M., Siller, R., Liang, K. X., Sullivan, G., et al. (2020). A method for differentiating human induced pluripotent stem cells toward functional cardiomyocytes in 96-well microplates. Sci Rep-uk 10, 18498. doi:10.1038/s41598-020-73656-2.

Belmonte, S., and Morad, M. (2008). Shear fluid-induced Ca2+ release and the role of mitochondria in rat cardiac myocytes. Ann Ny Acad Sci 1123, 58–63. doi:10.1196/annals.1420.007.

Birket, M. J., Orr, A. L., Gerencser, A. A., Madden, D. T., Vitelli, C., Swistowski, A., et al. (2011). A reduction in ATP demand and mitochondrial activity with neural differentiation of human embryonic stem cells. J Cell Sci 124, 348–58. doi:10.1242/jcs.072272.

Calvo, S. E., Clauser, K. R., and Mootha, V. K. (2016). MitoCarta2.0: an updated inventory of mammalian mitochondrial proteins. Nucleic Acids Res 44, D1251–7. doi:10.1093/nar/gkv1003.

Cho, Y. M., Kwon, S., Pak, Y. K., Seol, H. W., Choi, Y. M., Park, D. J., et al. (2006). Dynamic changes in mitochondrial biogenesis and antioxidant enzymes during the spontaneous differentiation of human embryonic stem cells. Biochem Bioph Res Co 348, 1472–8. doi:10.1016/j.bbrc.2006.08.020.

Cliff, T. S., Wu, T., Boward, B. R., Yin, A., Yin, H., Glushka, J. N., et al. (2017). MYC Controls Human Pluripotent Stem Cell Fate Decisions through Regulation of Metabolic Flux. Cell Stem Cell 21, 502–516.e9. doi:10.1016/j.stem.2017.08.018.

Cogliati, S., Enriquez, J. A., and Scorrano, L. (2016). Mitochondrial Cristae: Where Beauty Meets Functionality. Trends Biochem Sci 41, 261–273. doi:10.1016/j.tibs.2016.01.001.

Cogliati, S., Frezza, C., Soriano, M. E., Varanita, T., Quintana-Cabrera, R., Corrado, M., et al. (2013). Mitochondrial cristae shape determines respiratory chain supercomplexes assembly and respiratory efficiency. Cell 155, 160–71. doi:10.1016/j.cell.2013.08.032.

Dai, D. F., Danoviz, M. E., Wiczer, B., Laflamme, M. A., and Tian, R. (2017). Mitochondrial Maturation in Human Pluripotent Stem Cell Derived Cardiomyocytes. Stem Cells Int 2017, 5153625. doi:10.1155/2017/5153625.

Divakaruni, A. S., and Brand, M. D. (2011). The regulation and physiology of mitochondrial proton leak. Physiology 26, 192–205. doi:10.1152/physiol.00046.2010.

Dykens, J. A., and Stout, A. K. (2001). Assessment of mitochondrial membrane potential in situ using single potentiometric dyes and a novel fluorescence resonance energy transfer technique. 65, 285–309. doi:Doi 10.1016/S0091-679x(01)65018-0.

Ekstrand, M. I., Falkenberg, M., Rantanen, A., Park, C. B., Gaspari, M., Hultenby, K., et al. (2004). Mitochondrial transcription factor A regulates mtDNA copy number in mammals. Hum Mol Genet 13, 935–44. doi:10.1093/hmg/ddh109.

Floros, V. I., Pyle, A., Dietmann, S., Wei, W., Tang, W. C. W., Irie, N., et al. (2018). Segregation of mitochondrial DNA heteroplasmy through a developmental genetic bottleneck in human embryos. Nat Cell Biol 20, 144–151. doi:10.1038/s41556-017-0017-8.

Folmes, C. D., Dzeja, P. P., Nelson, T. J., and Terzic, A. (2012). Metabolic plasticity in stem cell homeostasis and differentiation. Cell Stem Cell 11, 596–606. doi:10.1016/j.stem.2012.10.002.

Friedman, C. E., Nguyen, Q., Lukowski, S. W., Helfer, A., Chiu, H. S., Miklas, J., et al. (2018). Single-Cell Transcriptomic Analysis of Cardiac Differentiation from Human PSCs Reveals HOPX-Dependent Cardiomyocyte Maturation. Cell Stem Cell 23, 586–598 e8. doi:10.1016/j.stem.2018.09.009.

Gaare, J. J., Nido, G. S., Sztromwasser, P., Knappskog, P. M., Dahl, O., Lund-Johansen, M., et al. (2018). Rare genetic variation in mitochondrial pathways influences the risk for Parkinson’s disease. Movement Disord 33, 1591–1600. doi:10.1002/mds.64.

Gillis, J., Mistry, M., and Pavlidis, P. (2010). Gene function analysis in complex data sets using ErmineJ. Nat Protoc 5, 1148–59. doi:10.1038/nprot.2010.78.

Gong, G., Song, M., Csordas, G., Kelly, D. P., Matkovich, S. J., and Dorn, G. W. (2015). Parkin-mediated mitophagy directs perinatal cardiac metabolic maturation in mice. Science 350, aad2459. doi:10.1126/science.aad2459.

Hom, J. R., Quintanilla, R. A., Hoffman, D. L., Bentley, K. L. de M., Molkentin, J. D., Sheu, S. S., et al. (2011). The permeability transition pore controls cardiac mitochondrial maturation and myocyte differentiation. Dev Cell 21, 469–78. doi:10.1016/j.devcel.2011.08.008.

John, J. C. S., Ramalho-Santos, J., Gray, H. L., Petrosko, P., Rawe, V. Y., Navara, C. S., et al. (2005). The expression of mitochondrial DNA transcription factors during early cardiomyocyte in vitro differentiation from human embryonic stem cells. Cloning Stem Cells 7, 141–53. doi:10.1089/clo.2005.7.141.

Kanehisa, M., and Goto, S. (2000). KEGG: kyoto encyclopedia of genes and genomes. Nucleic Acids Res 28, 27–30. doi:10.1093/nar/28.1.27.

Kaufman, B. A., Durisic, N., Mativetsky, J. M., Costantino, S., Hancock, M. A., Grutter, P., et al. (2007). The mitochondrial transcription factor TFAM coordinates the assembly of multiple DNA molecules into nucleoid-like structures. Mol Biol Cell 18, 3225–3236. doi:10.1091/mbc.e07-05-0404.

Kukat, C., and Larsson, N. G. (2013). mtDNA makes a U-turn for the mitochondrial nucleoid. Trends Cell Biol 23, 457–63. doi:10.1016/j.tcb.2013.04.009.

Larsson, N. G., Oldfors, A., Holme, E., and Clayton, D. A. (1994). Low levels of mitochondrial transcription factor A in mitochondrial DNA depletion. Biochem Bioph Res Co 200, 1374–81. doi:10.1006/bbrc.1994.1603.

Larsson, N. G., Wang, J. M., Wilhelmsson, H., Oldfors, A., Rustin, P., Lewandoski, M., et al. (1998). Mitochondrial transcription factor A is necessary for mtDNA maintenance and embryogenesis in mice. Nature Genetics 18, 231–236. doi:DOI 10.1038/ng0398-231.

Lees, J. G., Gardner, D. K., and Harvey, A. J. (2018). Mitochondrial and glycolytic remodeling during nascent neural differentiation of human pluripotent stem cells. Development 145, dev168997. doi:10.1242/dev.168997.

Liang, K. X., Kristiansen, C. K., Mostafavi, S., Vatne, G. H., Zantingh, G. A., Kianian, A., et al. (2020). Disease-specific phenotypes in iPSC-derived neural stem cells with POLG mutations. Embo Mol Med 12, e12146. doi:10.15252/emmm.202012146.

Locasale, J. W., and Cantley, L. C. (2011). Metabolic Flux and the Regulation of Mammalian Cell Growth. Cell Metab 14, 443–451. doi:10.1016/j.cmet.2011.07.014.

Love, M. I., Huber, W., and Anders, S. (2014). Moderated estimation of fold change and dispersion for RNA-seq data with DESeq2. Genome Biol 15, 550. doi:10.1186/s13059-014-0550-8.

Lukyanenko, V., Chikando, A., and Lederer, W. J. (2009). Mitochondria in cardiomyocyte Ca2+ signaling. Int J Biochem Cell Biology 41, 1957–71. doi:10.1016/j.biocel.2009.03.011.

Malinska, D., Kudin, A. P., Bejtka, M., and Kunz, W. S. (2012). Changes in mitochondrial reactive oxygen species synthesis during differentiation of skeletal muscle cells. Mitochondrion 12, 144–8. doi:10.1016/j.mito.2011.06.015.

Miettinen, T. P., and Björklund, M. (2016). Cellular Allometry of Mitochondrial Functionality Establishes the Optimal Cell Size. Dev Cell 39, 370–382. doi:10.1016/j.devcel.2016.09.004.

Patro, R., Duggal, G., Love, M. I., Irizarry, R. A., and Kingsford, C. (2017). Salmon provides fast and bias-aware quantification of transcript expression. Nat Methods 14, 417–419. doi:10.1038/nmeth.4197.

Perry, S. W., Norman, J. P., Barbieri, J., Brown, E. B., and Gelbard, H. A. (2011). Mitochondrial membrane potential probes and the proton gradient: a practical usage guide. Biotechniques 50, 98– 115. doi:10.2144/000113610.

Rossmann, M. P., Dubois, S. M., Agarwal, S., and Zon, L. I. (2021). Mitochondrial function in development and disease. Dis Model Mech 14. doi:10.1242/dmm.048912.

Rowe, I., and Boletta, A. (2013). Mitochondrial Transmembrane Potential (Ψm) Assay Using TMRM. Bio-protocol 3. doi:10.21769/bioprotoc.987.

Sercel, A. J., Carlson, N. M., Patananan, A. N., and Teitell, M. A. (2021). Mitochondrial DNA Dynamics in Reprogramming to Pluripotency. Trends Cell Biol. doi:10.1016/j.tcb.2020.12.009.

Skelton, R. J. P., Kamp, T. J., Elliott, D. A., and Ardehali, R. (2017). Biomarkers of Human Pluripotent Stem Cell-Derived Cardiac Lineages. Trends Mol Med 23, 651–668. doi:10.1016/j.molmed.2017.05.001.

Soneson, C., Love, M. I., and Robinson, M. D. (2015). Differential analyses for RNA-seq: transcript-level estimates improve gene-level inferences. F1000research 4, 1521. doi:10.12688/f1000research.7563.2.

Sturzu, A. C., and Wu, S. M. (2011). Developmental and regenerative biology of multipotent cardiovascular progenitor cells. Circ Res 108, 353–64. doi:10.1161/circresaha.110.227066.

Taylor, K. D., and Pikó, L. (1995). Mitochondrial biogenesis in early mouse embryos: Expression of the mRNAs for subunits IV, Vb, and VIIc of cytochrome c oxidase and subunit 9 (P1) of H+-ATP synthase. Mol Reprod Dev 40, 29–35. doi:10.1002/mrd.1080400105.

Teixeira, F. K., Sanchez, C. G., Hurd, T. R., Seifert, J. R., Czech, B., Preall, J. B., et al. (2015). ATP synthase promotes germ cell differentiation independent of oxidative phosphorylation. Nat Cell Biol 17, 689–96. doi:10.1038/ncb3165.

Tzoulis, C., Tran, G. T., Schwarzlmuller, T., Specht, K., Haugarvoll, K., Balafkan, N., et al. (2013). Severe nigrostriatal degeneration without clinical parkinsonism in patients with polymerase gamma mutations. Brain 136, 2393–2404. doi:10.1093/brain/awt103.

Varum, S., Rodrigues, A. S., Moura, M. B., Momcilovic, O., Easley, C. A., Ramalho-Santos, J., et al. (2011). Energy Metabolism in Human Pluripotent Stem Cells and Their Differentiated Counterparts. Plos One 6, e20914. doi:10.1371/journal.pone.0020914.

Vliet, P. V., Wu, S. M., Zaffran, S., and Puceat, M. (2012). Early cardiac development: a view from stem cells to embryos. Cardiovasc Res 96, 352–62. doi:10.1093/cvr/cvs270.

Wanet, A., Arnould, T., Najimi, M., and Renard, P. (2015). Connecting Mitochondria, Metabolism, and Stem Cell Fate. Stem Cells Dev 24, 1957–1971. doi:10.1089/scd.2015.0117.

Wanet, A., Remacle, N., Najar, M., Sokal, E., Arnould, T., Najimi, M., et al. (2014). Mitochondrial remodeling in hepatic differentiation and dedifferentiation. Int J Biochem Cell Biology 54, 174–85. doi:10.1016/j.biocel.2014.07.015.

Zhang, J., Nuebel, E., Daley, G. Q., Koehler, C. M., and Teitell, M. A. (2012a). Metabolic Regulation in Pluripotent Stem Cells during Reprogramming and Self-Renewal. Cell Stem Cell 11, 589–595. doi:10.1016/j.stem.2012.10.005.

Zhang, J., Nuebel, E., Wisidagama, D. R., Setoguchi, K., Hong, J. S., Horn, C. M. V., et al. (2012b). Measuring energy metabolism in cultured cells, including human pluripotent stem cells and differentiated cells. Nat Protoc 7, 1068–85. doi:10.1038/nprot.2012.048.

Zheng, X., Boyer, L., Jin, M., Mertens, J., Kim, Y., Ma, L., et al. (2016). Metabolic reprogramming during neuronal differentiation from aerobic glycolysis to neuronal oxidative phosphorylation. Elife 5, e13374. doi:10.7554/elife.13374.

Zhu, H., Scharnhorst, K. S., Stieg, A. Z., Gimzewski, J. K., Minami, I., Nakatsuji, N., et al. (2017). Two dimensional electrophysiological characterization of human pluripotent stem cell-derived cardiomyocyte system. Sci Rep-uk 7, 43210. doi:10.1038/srep43210.

Zwi, L., Caspi, O., Arbel, G., Huber, I., Gepstein, A., Park, I. H., et al. (2009). Cardiomyocyte differentiation of human induced pluripotent stem cells. Circulation 120, 1513–23. doi:10.1161/circulationaha.109.868885.

